# A topological approach to DNA similarity analysis from 5-dimensional representation

**DOI:** 10.1101/2021.03.10.434824

**Authors:** Dong Quan Ngoc Nguyen, Phuong Dong Tan Le, Ziqing Hu, Lizhen Lin

**Affiliations:** Department of Applied and Computational Mathematics and Statistics, University of Notre Dame, Notre Dame, IN 46556, USA; Department of Applied Mathematics, University of Waterloo, Waterloo, Ontario, Canada N2L 3G1

## Abstract

In this paper, we propose another topological approach for DNA similarity analysis. For each DNA sequence, we transform it into a collection of vectors in 5-dimensional space in which all nucleotides of the same type, say A, C, G, T are on the same line in this 5D space. Based on this special geometric property, we combine this representation with tools in persistent homology to obtain only zeroth persistence diagrams as a topological representation of DNA sequences. Similarities between DNA sequences are signified via how close the representing zeroth persistence diagrams of the DNA sequences are, based on the Wasserstein distance of order zero, which provides a new method for analyzing similarities between DNA sequences. We test our methods on the datasets of Human rhinovirus (HRV) and Influenza A virus.

## 1 Introduction

There are two main approaches to the problem of comparing DNA sequences: alignment methods and alignment-free methods. In alignment-based methods, the classical multiple sequence alignment (MSA) method is widely used. Although the MSA method has the highest accuracy among DNA similarity analysis methods, time complexity becomes too large for a large dataset of long DNA sequences. In alignment-free methods, several methods have been proposed to reduce time complexity while maintaining a high accuracy in comparing DNA sequences. In general, an alignment-free method based on graphical representations of DNA sequences consists of two step, the first of which embed each DNA sequence into an Euclidean space, and the second is to compute the similarity matrix based on certain distance-based methods such as Euclidean distances or correlation angles to realize differences or similarities among DNA sequences. For papers using graphical representation for DNA similarity analysis, see, for example, [1, 2, 3, 4, 5, 6, 7, 8, 9, 10, 11, 12, 13, 14, 15, 16, 17, 18, 19, 20].

In a recent work, Nguyen, Le, Xing, and Lin [21] proposed a different alignment-free approach which combines geometry with topology of DNA sequences. In [21], a new 4D representation of DNA sequences was introduced using a chaos in the four-dimensional Euclidean space ℝ^4^. Instead of computing similarities/dissimilarities between DNA sequences based on Euclidean distances or correlation angles as in other work, Nguyen, Le, Xing, and Lin [21] proposed to explore topological properties of the sets of 4-dimensional vectors that represent DNA sequences in order to obtain intrinsic geometrical and topological structures of DNA sequences for similarity analysis. The main mathematical tools used in [21] are chaos in high dimensions, and persistent homology which has recently gained an important role in topological data analysis and related areas. In this paper, we follow a similar strategy of using persistent homology as in [21] for DNA similarity analysis, but we use a different geometric 5-dimensional representation that is based on that of Liao, Li, and Zhu [22]. In [21], Nguyen, Le, Xing, and Lin used chaos representation in the 4-dimensional space ℝ^4^ under which persistence diagrams of DNA sequences will contain nontrivial higher-dimensional homology groups, which in turn requires more powerful computation power for performing similarity analysis between DNA sequences, based on Wasserstein distances of order at least one. The main difference of our present method from [21] is that under our geometric 5D representation of DNA sequences, persistent homology only contains the zeroth homology group, which in turn greatly simplifies time complexity for analyzing similarities between DNA sequences, based on the Wasserstein distance of order zero. Furthermore the zeroth persistence diagrams of DNA sequences based on the 5D representation of DNA sequences provide a *simplest possible* topological visualization of DNA sequences which in many cases provides a quick assessment of similarities/dissimilarities between DNA sequences.

## 2 Method

Our method consists of two steps. In the first step, we transform each DNA sequence into the 5-dimensional Euclidean space ℝ^5^ so that each DNA sequence of length *n* can be represented by a collection of *n* vectors in ℝ^5^. In the second step, we compute persistent homology for each such collection of vectors to obtain the persistence diagrams of DNA sequences which contain intrinsic topological information of each DNA sequence. The 5D representation that we use only leads to the nontrivial zeroth persistence diagrams of DNA sequences which provide a simplest possible topological visualization of DNA sequences. It is known that the collection of zeroth persistence diagrams is a metric under the Wasserstein distance of order zero. In order to compare similarities between DNA sequences, we compute the similarity/dissimilarity matrix using the Wasserstein distance.

### 2.1 5-dimensional representation of DNA sequences

In this subsection, we introduce a map that transforms each DNA sequence of length *n* into a collection of *n* vectors in ℝ^5^. Our construction is based on that in Liao, Li, and Zhu [22] with slight modification. Let *α* = *a*_1_*a*_2_ · · · *a_n_* be a DNA sequence of length *n*, where the *a_i_* denotes one of 4 nucleotide bases A, C, G, T. For each 1 ≤ *i* ≤ *n*, set

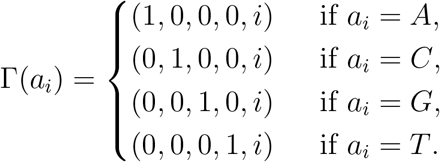

In [22], Liao, Li, and Zhu maps *a_i_* = *G* to (0, 1, 0, 0, *i*), and *a_i_* = *C* to (0, 0, 1, 0, *i*) which is a permutation of the above construction.

Figure **??** illustrates the 3D projection of the 5-dimensional representation of the segment from nucleotides 3660 to 3669 of the DNA sequence of Human Rhinovirus of type A-HRV-65 (whose accession number is FJ445147) (see [25]).

### 2.2 Persistent homology and persistent diagrams

We briefly recall the notion of persistent homology and persistence diagrams that will apply for analyzing DNA sequences. The reader is referred to [23] for a detailed reference about persistent homology and persistence diagrams. For a given nonnegative integer *k* ≥ 0 and a collection of *k* + 1 points *u*_0_, …, *u_k_* in ℝ^*k*+1^. One can create a **convex hull** of this collection in ℝ^*k*+1^ by including all convex combinations of these points of the form 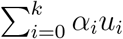, where the *α_i_* are between 0 and 1 such that 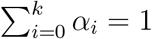. We call such convex hull the *k*-simplex generated by the points *u*_0_*, … , u_k_*, and denote it by [*u*_0_*, … , u_k_*]. For a collection of simplexes in ℝ^*k*+1^, say Δ. We call Δ a **simplicial complex** if whenever *σ* is a simplex in Δ, all *d*-simplexes contained in *σ* are also contained in Δ. For such a geometric object Δ in ℝ^*n*^, based on algebraic topology, there exists, for each *j* ≥ 0, an algebraic structure called the *j***-th homology group of** Δ, denoted by 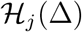 which behaves in a similar way as a vector space over ℝ. There is an analogue of dimensions of vector spaces over ℝ in the setting of homology groups 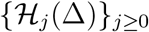 that we call the **rank of** 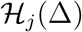, a positive integer, which signifies important geometric properties of Δ. For example, the rank of 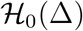 equals the *number of connected components of* Δ in ℝ^*n*^, and the rank of 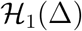 denotes the *number of* 1*-dimensional holes* of Δ.

Let *X* be a finite set of points, say *a*_1_, …, *a_m_* in ℝ^*n*^, and let *d* denote the standard Euclidean distance in ℝ^*n*^. For each *ϵ* ≥ 0, set

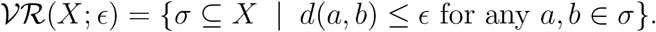

One can verify that 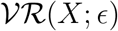 is a simplicial complex called the *ϵ***-Vietoris-Rips complex of** *X*. Let *ϵ*_0_ = −∞ < 0 ≤ *ϵ*_1_ ≤ · · · ≤ *ϵ_h_* ≤ · · · be an increasing sequence of nonnegative real numbers. One can form a sequence of simplicial complexes 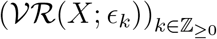, and one obtains a **filtration of the form**

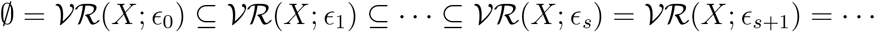

which will stabilize at some point *ϵ_s_*. For each 0 ≤ *p* ≤ *q* ≤ *s*, the embedding 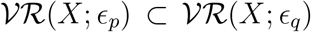 induces a sequence of natural maps 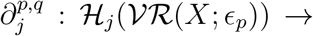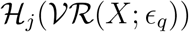. For each *j* ≥ 0, the *j***-th persistent homology groups** 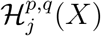 **of** *X* are the images of 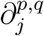 which are 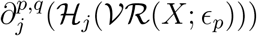.

Each element *γ* in 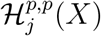 is called a *j***-topological feature of** *X*. The *j***-th persistence diagram of** *X* is a set of points ((*b, d*) | 0 ≤ *b* < *d*} ⊂ ℝ^2^, where each point (*b, d*) signifies the **birth and death times of a** *j***-topological feature** *γ* of *X*, i.e., *b* is the radius in which *γ* first appears in 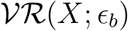 and *d* is the radius in which *γ* gets filled in with a lower dimensional simplex. We denote by 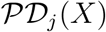 the *j***-th persistence diagram of** *X*. In our methods, it suffices to consider only the 0-th persistence diagrams, which correspond to topological features of connectedness of *X*.

Let *X, Y* be two finite sets of points in ℝ^*n*^. In order to compare topological features of *X, Y* in our methods, we consider the **Wasserstein distance of degree** 0 between 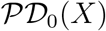 and 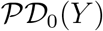, i.e.,

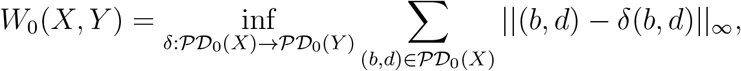

where || · ||_∞_ denotes the *L*_∞_-distance between two points in ℝ^2^.

### 2.3 Proposed Method

Our proposed method for reconstructing a phylogenetic tree of DNA sequences is described in the following algorithm:

(0) (Input) A collection of *n* DNA sequences *α*_1_*, …, α_n_*.
(1) Construct the 5-dimensional geometric representation of each DNA sequence *α_i_* from Subsection 2.1 to obtain a finite set of points 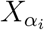 in ℝ^5^.
(2) Compute the 1st persistence diagrams of the 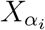 to obtain the sets of 0-th persistence diagrams 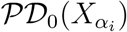 in ℝ^2^, using the notions in Section 2.2. We use Python packages from https://pypi.org/project/persim/ to compute persistence diagrams and Wasserstein distances. Note that using the 5-dimensional representation in Subsection 2.1, all *j*-th persistence diagrams with *j* ≥ 1 are empty, which leads to a very simple way to compare DNA sequences based on the 0-th persistence diagrams.
(3) Compute the distance matrix of dimensions *n* × *n* whose (*i, j*)-entry is the Wasserstein distance 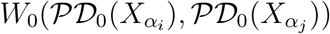.
(4) (Ouput) Construct the phylogenetic tree of the DNA sequences from the distance matrix in Step 3, using UPGMA algorithm (see [24]).

## 3 Results

In this section, we apply our method described in Section 2 to analyzing three datasets: Human rhinovirus, Influenza A virus, and Human Papillomavirus (HPV). In the HRV dataset, there are 113 DNA sequences whose accession numbers on the GenBank (see https://www.ncbi.nlm.nih.gov/genbank/) can be obtained from Appendix A (Supplementary data) of Hoang, Yin, and Yau [29]. The Influenza dataset has 38 DNA sequences whose accession numbers on the GenBank can also be found from the same Appendix of [29].

### 3.1 Human rhinovirus (HRV)

HRV is the most common viral infectious agent in humans, and is the main cause of the common cold. In [25], using multiple sequence alignment, Palmenberg et al. [25] correctly classified the complete HRV genomes into three genetically distinct groups within the genus *Enterovirus* (HEV) and the family *Picornaviridae*. The dataset used in [25] consists of three groups HRV-A, HRV-B, HRV-C including 113 genomes, and three outgroup sequences HEV-C. The time complexity was very high because of the use of multiple sequence alignment. In this paper, we use the same dataset to test our method.

From the phylogenetic tree of HPV genomes based on our method in Figure 2, we find that except some genomes of type HRV-C inaccurately grouped together with type HRV-A, and some genomes of type HRV-A being away from the main branch of type HRV-A, all other genomes are correctly classified into their corresponding types.

**Figure 1:**
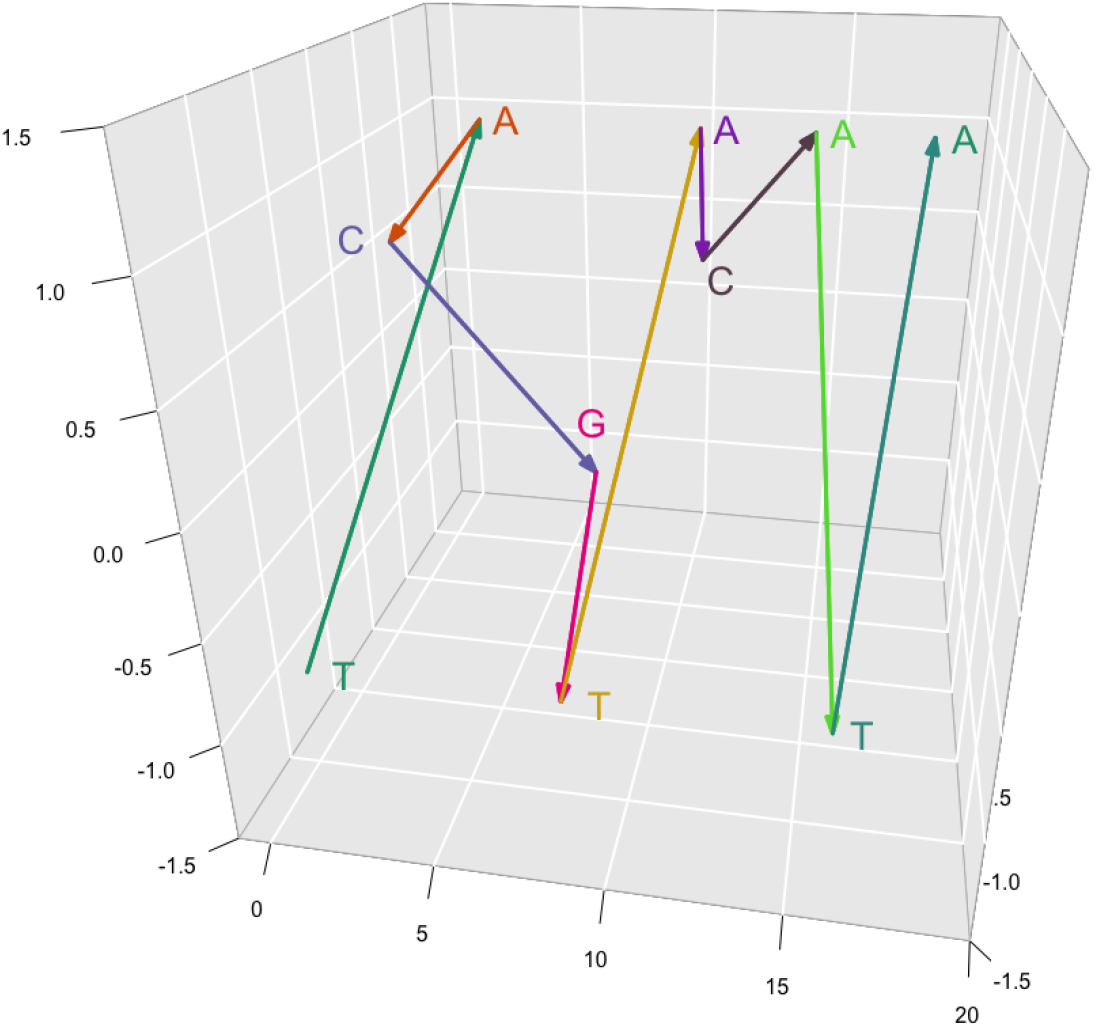
3D projection of the 5D representation of the DNA sequence of Human Rhi-novirus of type A-HRV-65

**Figure 2:**
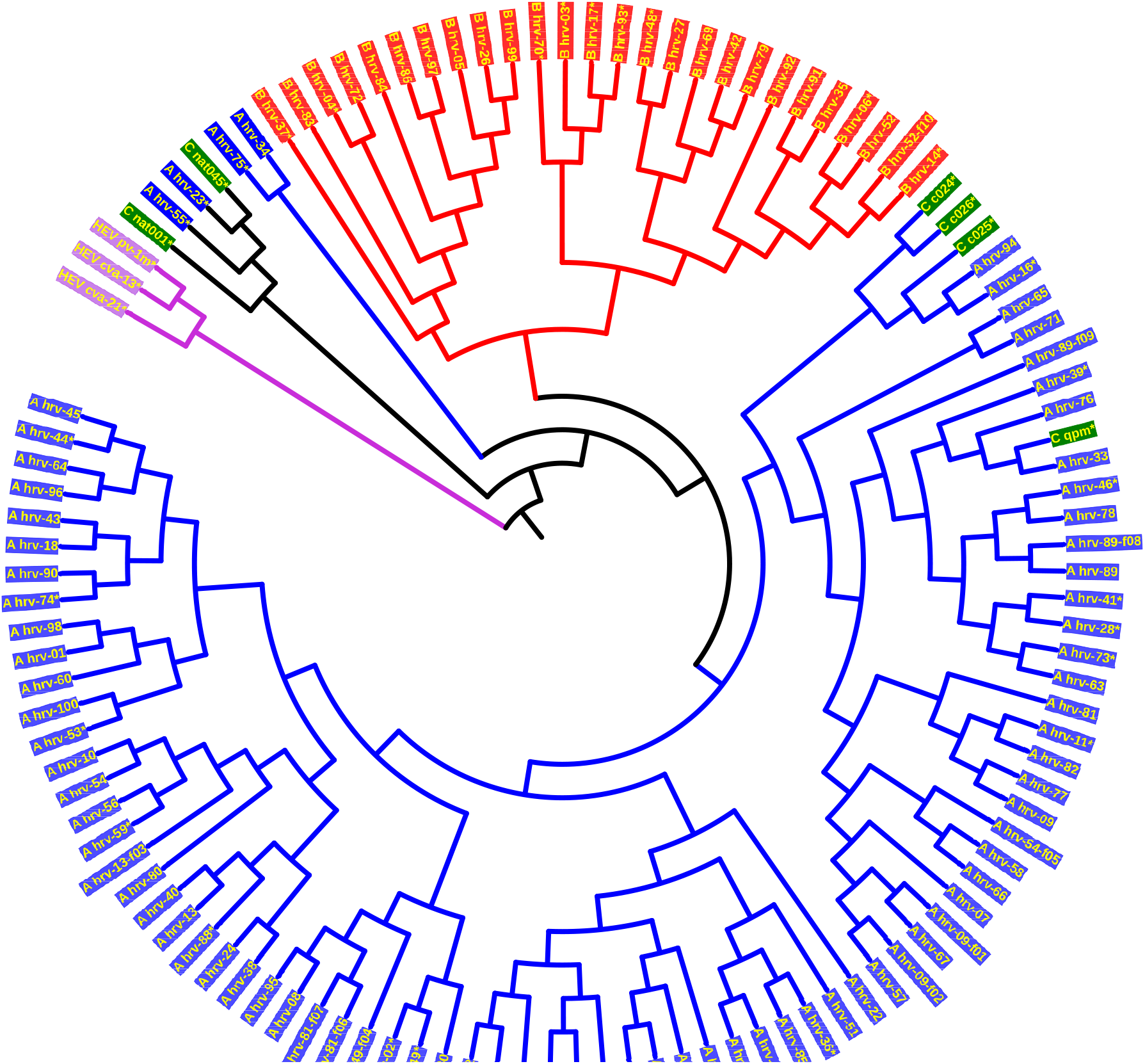
Phylogenetic tree of 113 HRV genomes of 4 genotypes

### 3.2 Influenza

Influenza A viruses are very dangerous because they have a wide range of hosts including birds, horses, swine, and humans. These viruses have been a serious health threat to humans and animals (see [26]), and are known to have high degree of genetic and antigenic variability (see [27, 28]). Some subtypes of Influenza A viruses are very dangerous, and lethal including H1N1, H2N2, H5N1, H7N3, and H7N9. We apply our method on the dataset consisting of 38 Influenza A virus genomes whose accession numbers in GenBank can be found in the Appendix A (Supplementary data) of [29]. From Figure 4, we find that except A/American black duck/NB/2538/2007-H7N3, A/chicken/British Columbia/GSC human B/04-H7N3, A/turkey/Minnesota/1/1988-H7N9 inaccurately misplaced in H2N2 group, all other Influenza A genomes are clustered correctly into their types.

In addition, our proposed method allows one to visually inspect differences between Influenza A virus genomes in the *simplest possible way* of persistent homology. For example, Figure 3 illustrates identical 0-th persistence diagrams of *A/mallard/Maryland/352/2002-H1N1* and *A/mallard/Maryland/26/2003-H1N1* whose highly identical visualization shows that they should belong in the same branch as indicated by Figure 4. Furthermore the 0-th persistence diagram *A/mallard/Maryland/352/2002-H1N1* shows that the geometric shape of its DNA sequence has exactly two connected components.

**Figure 3:**
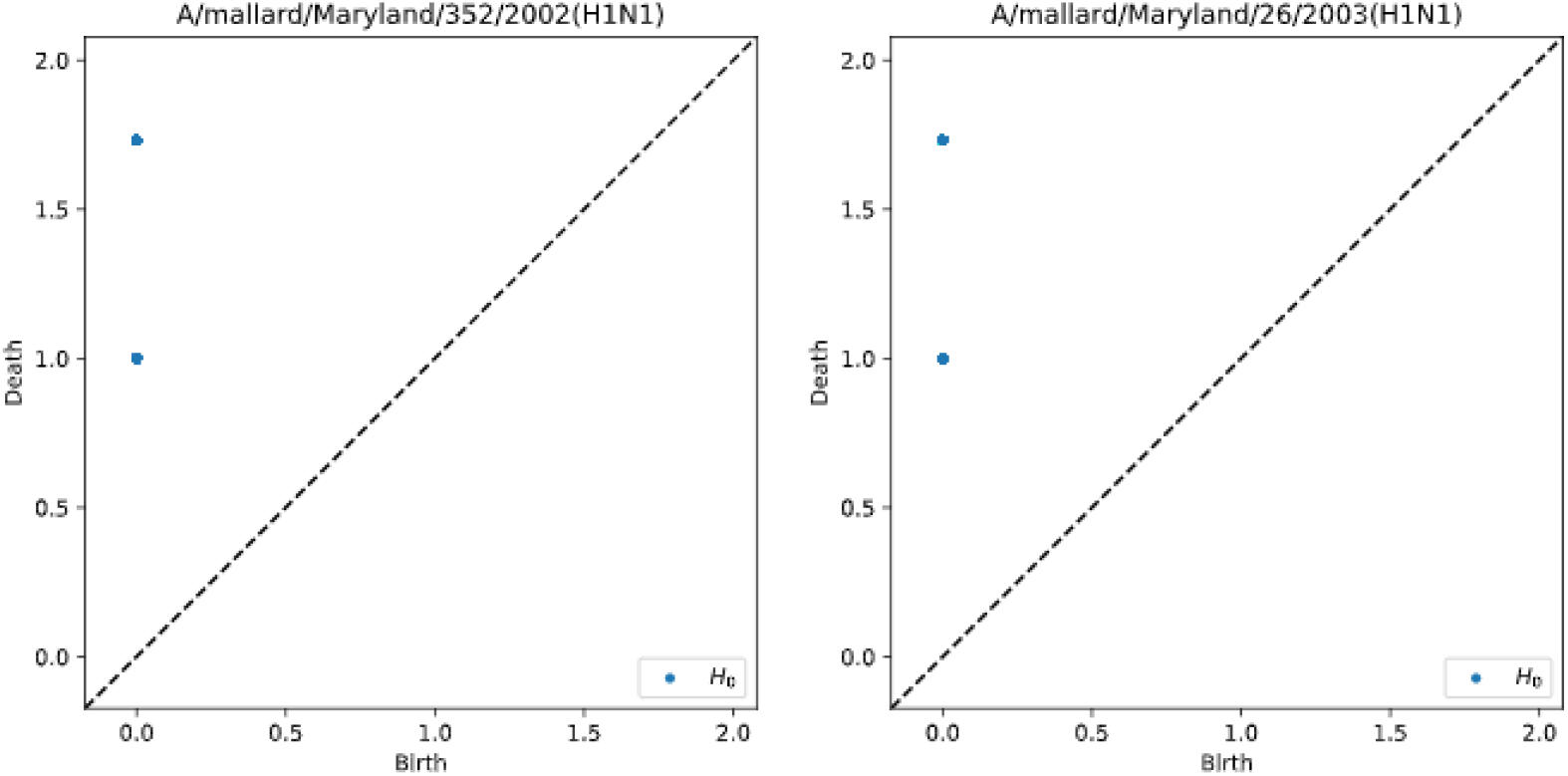
0-th persistence diagrams of Influenza A virus genomes

**Figure 4:**
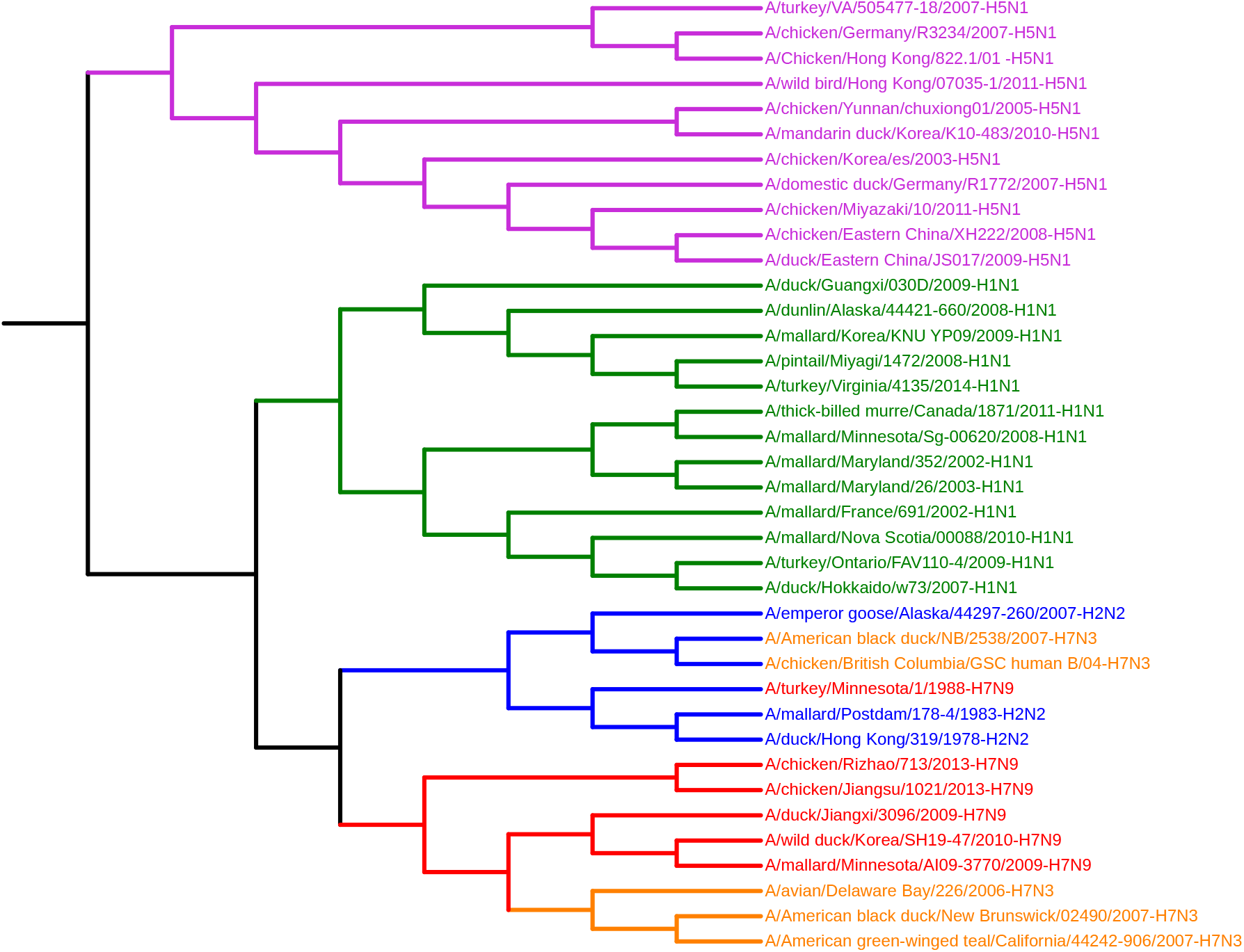
Phylogenetic tree of 38 Influenza A virus genomes

## Notes

### Competing Interest Statement

The authors have declared no competing interest.

